# Amortized Hypothesis Generation

**DOI:** 10.1101/137190

**Authors:** Ishita Dasgupta, Eric Schulz, Noah D. Goodman, Samuel J. Gershman

## Abstract

Bayesian models of cognition posit that people compute probability distributions over hypotheses, possibly by constructing a sample-based approximation. Since people encounter many closely related distributions, a computationally efficient strategy is to selectively reuse computations – either the samples themselves or some summary statistic. We refer to these reuse strategies as *amortized inference*. In two experiments, we present evidence consistent with amortization. When sequentially answering two related queries about natural scenes, we show that answers to the second query vary systematically depending on the structure of the first query. Using a cognitive load manipulation, we find evidence that people cache summary statistics rather than raw sample sets. These results enrich our notions of how the brain approximates probabilistic inference.

## Introduction

Many theories of probabilistic reasoning assume that people are equipped with a general-purpose inference engine that can be used to answer arbitrary queries for a wide variety of probabilistic models (Griffiths et al., 2012). The flexibility and power of a general-purpose inference engine trades off against its computational efficiency: by avoiding any assumptions about the query distribution, the inference engine relinquishes the opportunity to reuse computations across queries. Conversely, an inference engine may gain efficiency by incurring some amount of bias due to reuse—a strategy we refer to as *amortized inference* (Stuhlmüller et al., 2013; Gershman & Goodman, 2014). We propose that people employ some form of this strategy, flexibly reusing past inferences in order to efficiently answer new but related queries.

The experiments reported in this paper explore amortization in sets of related queries that involve probabilistic inference over a very large space of possibilities. These possibilities are not all explicitly provided and have to be generated by the participant in order to carry out the inference. We frame amortization as the reuse of hypotheses that have already been generated in response to previous queries. We model the process of hypothesis generation with a stochastic sampling mechanism (Lieder et al., 2012; Dasgupta et al., 2016). One way to implement amortization in this framework is to directly reuse samples across different queries. Alternatively, amortization could be implemented by reusing some summary statistic compiled from previous samples. One goal of our experiments is to tease apart these different mechanistic assumptions. The basic logic of our experiments is to hold one query fixed while manipulating an earlier query, allowing us to interrogate reuse of computations across queries.

### Stochastic hypothesis generation

For complex hypothesis spaces, exact probabilistic inference is typically intractable. Several lines of evidence point to the idea that humans approximate inference by constructing a Monte Carlo approximation of the posterior distribution (see Griffiths et al., 2012; Sanborn & Chater, 2016, for a review). This “sampling hypothesis” can be realized algorithmically in a number of ways. Recent studies have shown how a number of apparent probabilistic fallacies can be understood as a consequence of resource-bounded sampling using a *Markov chain Monte Carlo* (MCMC) algorithm (Lieder et al., 2012; Dasgupta et al., 2016). Because we build on those studies in this paper, we briefly describe the theoretical framework.

A Monte Carlo approximation uses a set of *N* samples *h*_1_*, …, h*_*N*_, drawn from a hypothesis space ℋ, to approximate a target distribution. In our case, the target is the posterior distribution over hypotheses given data *d*, *P*(*h|d*) ∝ *P*(*d|h*)*P*(*h*). The Monte Carlo approximation is defined by:

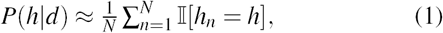

where 𝕝[·] = 1 if its argument is true (0 otherwise). MCMC algorithms generate samples by simulating a Markov chain whose stationary distribution is the posterior (MacKay, 2003). This approach is asymptotically exact (the approximation converges to the posterior as the number of samples approaches infinity) but under time or resource constraints only a small number of samples may be generated. Although this gives rise to systematic deviations from exact inference, it may in fact be the computationally rational sampling policy (Lieder et al., 2012; Vul et al., 2014; Gershman et al., 2015).

In our prior research (Dasgupta et al., 2016), we applied this model to a scene inference domain, using a database of natural object co-occurrence statistics compiled by Greene (2013). The task facing subjects in our experiments was to judge the probability of a particular set of latent objects in a scene conditional on observing another object (the cue). By manipulating the framing of the query, we showed that subjects gave different answers to formally equivalent queries. Specifically, by partially unpacking the queried object set (where fully unpacking an object set means to present it explicitly as a union of each of its member objects) into a small set of exemplars and a ‘catch-all’ hypothesis (e.g., “what is the probability that there is a book, a box, or any other object beginning with B?”), we found that subjects judged the probability to be higher when the unpacked exemplars were typical (a “subadditivity” effect; cf. Tversky & Koehler, 1994) and lower when the unpacked exemplars were atypical (a “superadditivity” effect; cf. Sloman et al., 2004) compared to when the query is presented without any unpacking. To give a concrete example, in the presence of say a ‘table’, the typically unpacked query “what is the probability that there is also a chair, a curtain, or any other object beginning with C?” generates higher probability estimates, and the atypically unpacked query “what is the probability that there is also a cow, a canoe, or any other object beginning with C?” generates lower probability estimates, when compared to the packed query “what is the probability that there is also another object beginning with C?”.

These effects could be accounted for by the MCMC model under the assumption that the unpacked exemplar(s) initialize the Markov chain(s) that form the sample set. Because the initialization of the Markov chain transiently determines its future trajectory, initializing with typical examples causes the chain to tarry in the high probability region of the queried object set, thereby increasing its judged probability (subadditivity). However, initializing with atypical examples causes the chain to get easily derailed into regions outside the queried object set and thus generate more hypotheses that are not in the queried object set. This decreases the judged probability of the queried object set (superadditivity). The strength of these effects diminishes with the number of samples, as the chain approaches its stationary distribution (which is the same for all conditions). Accordingly, response time and cognitive resource availability modulate the effect size (Dasgupta et al., 2016).

### Amortized inference in hypothesis generation

We will consider two simple amortization schemes within the MCMC framework described above.^1^ In *sample reuse*, people may simply add samples generated in response to one query (*Q*1) to the sample set for another query (*Q*2). So if *N*_1_ samples were generated in response to *Q*1, and *N*_2_ new samples are generated in response to *Q*2, the response to *Q*2 is given by a calculation carried out over all *N*_1_ + *N*_2_ equally weighted samples. Under this scheme, all the computations carried out for *Q*1 are available for flexible reuse in the computation for *Q*2. In *summary reuse*, people may reuse a summary statistic computed from *Q*1. This strategy is only applicable to problems where the answer to *Q*2 can be expressed as the composition of the answer to *Q*1, and an additional simpler computation. To make this viable in our experiments, *Q*2 queries a hypothesis space that is the union between the hypothesis space queried in *Q*1 and a disjoint hypothesis space. For example if *Q*1 is “What is the probability that there is an object starting with a C in the scene?”, *Q*2 could be “What is the probability that there is an object starting with a C or an R in the scene?”. In this case, the *N*_1_ samples generated in response to *Q*1 are summarized into one probability (“the probability of an object starting with C”), *N*_2_ new samples are generated in response to a simpler query (“the probability of an object starting with R”), and these two numbers are then composed (in this case simply added) to give the final estimate for *Q*2 (“the probability of an object starting with C or R”). Under this scheme, only the final product of the computation carried out for *Q*1 is reused in the calculations for *Q*2. These two models are the two extremes between very flexible and very rigid reuse; intermediates are of course possible.

Sample-based and summary-based amortization schemes make different predictions about how subadditivity and superadditivity change as a function of the sample size (Figure 1). Increasing the sample size for *Q*1 amplifies the effects for *Q*2 under a sample-based scheme, because this leads to more *Q*1 samples being reused for *Q*2. This effect can be counteracted by increasing the sample size for *Q*2, which pushes the effects down (the effects go to 0 as the sample size for *Q*2 tends to infinity, since the Markov chain will converge to the same posterior for all conditions). Under a summary-based scheme, increasing the sample size for *Q*1 will actually *diminish* the effects for *Q*2, because the bias from *Q*1 is strongest when the chain is close to its starting point. In other words, the subadditivity and superadditivity effects for *Q*2 derive from the same effects in *Q*1, which themselves are primarily driven by the initialization (see Dasgupta et al., 2016). In Experiment 2, we test these different predictions by placing people under cognitive load during either *Q*1 or *Q*2, a manipulation that is thought to reduce the number of samples (Dasgupta et al., 2016; Thaker et al., 2017).

**Figure 1:**
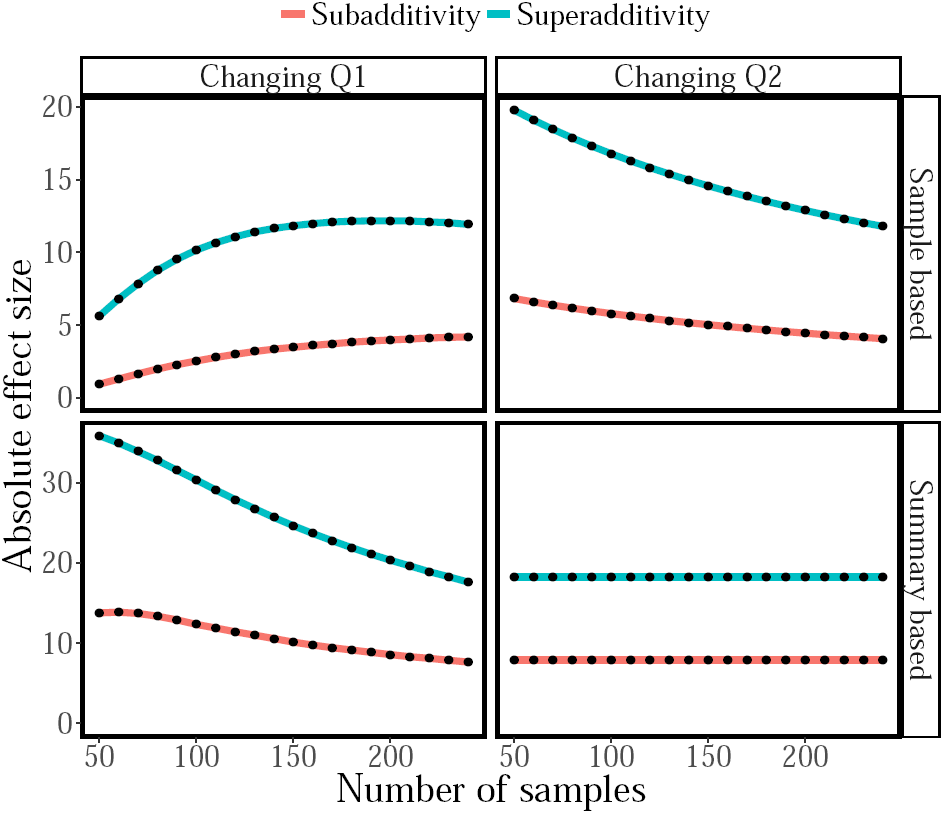
Simulation of subadditivity and superadditivity effects under sample-based (top) and summary-based (bottom) amortization schemes. In all panels, the y-axis represents the effect size for *Q*2. Left panels show the effects of changing the sample size for *Q*1; right panels show the effects of changing the sample size for *Q*2. When, sample size for *Q*2 is changed, sample size for *Q*1 is held fixed at 230, and vice versa.

## Experiment 1

Our first experiment pursued a basic carryover effect from one query (*Q*1) to the next (*Q*2). We assessed two putative signatures of sampling—subadditivity and superadditivity—for a fixed *Q*2 while changing the structure of *Q*1. Specifically, we compared three conditions that differed only in how *Q*1 was framed: packed, unpacked-typical, and unpacked-atypical. In order to encourage amortization, we specified *Q*2 as a union of the hypothesis space queried by *Q*1 and another hypothesis space—i.e., a disjunctive query. The design is summarized in Table 1.

**Table 1:** Experimental stimuli and queries.

| Cue | Q1 | Q1-Typical | Q1-Atypical | Q2 |
| --- | --- | --- | --- | --- |
| Table | C | chair,<br>computer<br>curtain | cannon,<br>cow,<br>canoe | C or R |
| Tele-<br>phone | D | display case,<br>dresser,<br>desk | drinking fountain,<br>dryer,<br>dome | D or L |
| Rug | B | book,<br>bouquet,<br>bed | bird,<br>buffalo,<br>bicycle | B or S |
| Chair | P | painting,<br>plant,<br>printer | porch,<br>pie,<br>platform | P or A |
| Sink | T | table,<br>towel,<br>toilet | trumpet,<br>toll gate,<br>trunk | T or E |
| Lamp | W | window,<br>wardrobe,<br>wine rack | wheelbarrow,<br>water fountain,<br>windmill | W or F |

### Participants

84 participants (53 males, mean age=32.61, SD=8.79) were recruited via Amazon’s Mechanical Turk and received $0.5 for their participation plus an additional bonus of $0.1 for every on-time response.

### Procedure

Participants were asked to imagine playing a game in which their friend sees a photo and then mentions one particular object present in the photo (the cued object). The participant is then queried about the probability that another class of objects (e.g., “objects beginning with the letter B”) is also present in the photo.

Each participant completed 6 trials, where the stimuli on every trial corresponded to the rows in Table 1. On each trial, participants first answered *Q*1 given the cued object, using a slider bar to report the conditional probability (Figure 2). The *Q*1 framing (typical-unpacked, atypical-unpacked or packed) was chosen randomly. Participants then completed the same procedure for *Q*2, conditional on the same cued object. The framing for *Q*2 was always packed.

**Figure 2:**
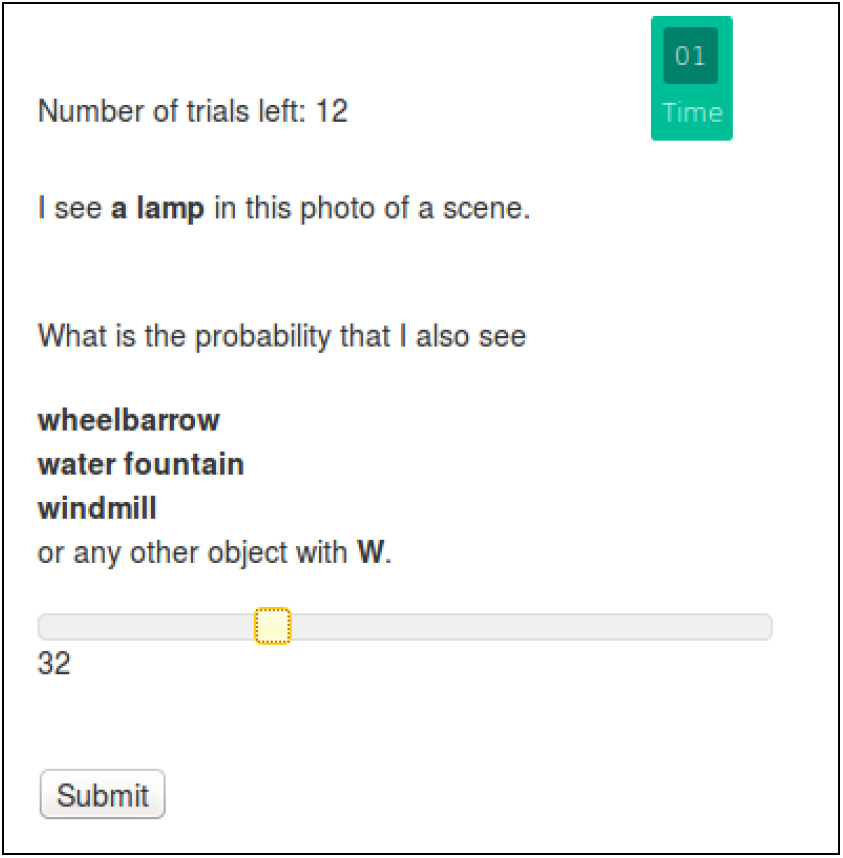
Experimental setup. Participants were asked to estimate the conditional probability using a slider bar within a 20-second time limit.

### Results

Consistent with our prior studies (Dasgupta et al., 2016), we find both subadditivity and superadditivity effects for *Q*1, depending on the unpacking (Figure 3): probability judgments were higher for unpacked-typical queries than for packed queries (a subadditivity effect; *t*(77) = 4.029*, p <* 0.001) and lower for unpacked-atypical than for packed queries (a superadditivity effect; *t*(77) = *-*6.4419*, p <* 0.001)

**Figure 3:**
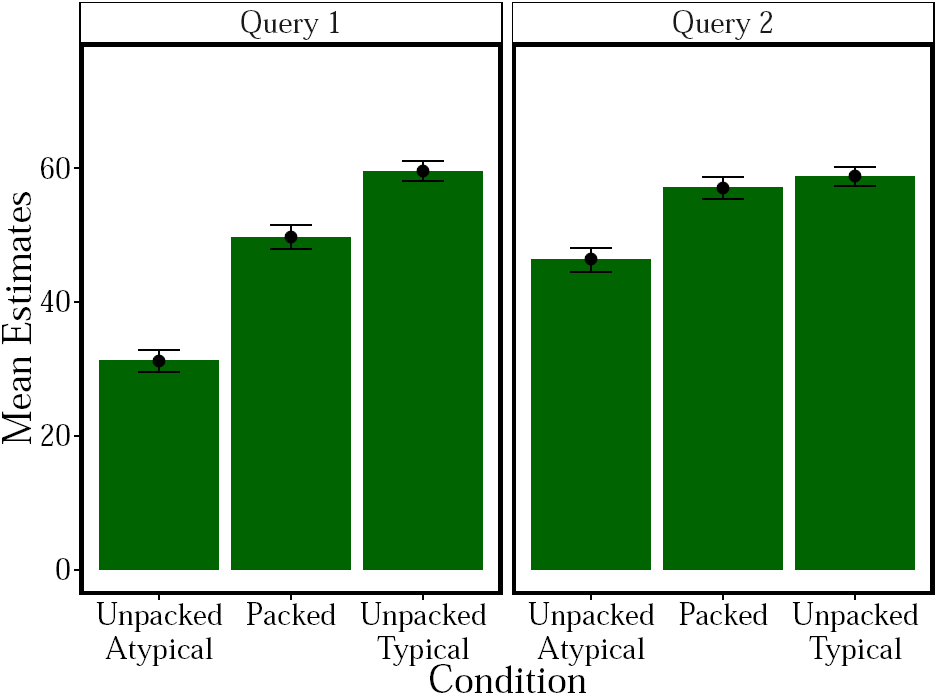
Experiment 1 results. Mean probability estimates by condition. Error bars represent the standard error of the mean.

Crucially, we also saw effects of *Q*1 unpacking on response to *Q*2, even though these queries were all presented as packed hypotheses. In particular, estimates for *Q*2 were lower when *Q*1 was unpacked to atypical exemplars (*t*(77) = −5.0789, *p <* 0.001), demonstrating a superadditivity effect that carried over from one query to another. We did not find an analogous carry-over effect for subadditivity (*t*(77) = 0.72, *p >* 0.4), possibly due to the subadditivity effect “washing out” more quickly (i.e. with fewer samples) than superadditivity, at least within this domain (see Dasgupta et al., 2016).

Next, we explored whether responses to *Q*1 predicted trialby-trial variation in responses to *Q*2. As shown in Figure 4, we found significant positive correlations between the two queries in all conditions when aggregating across participants (*p <* 0.01). The same conclusion can be drawn from analyzing correlations within participants and then testing the average correlation against 0 (average correlation: *r* = 0.55, *p <* 0.01). Moreover, the within-participant effect size (the response difference between the unpacked conditions and the packed conditions) for *Q*1 was correlated with responses to *Q*2 for both atypical (*r* = 0.35, *p <* 0.01) and typical (*r* = 0.21, *p <* 0.05) unpacking conditions. This means that even though the subadditive condition did not significantly differ from the unpacked condition for *Q*2 overall, participants who showed greater subadditivity or superadditivity for *Q*1 also showed correspondingly greater effects for *Q*2.

**Figure 4:**
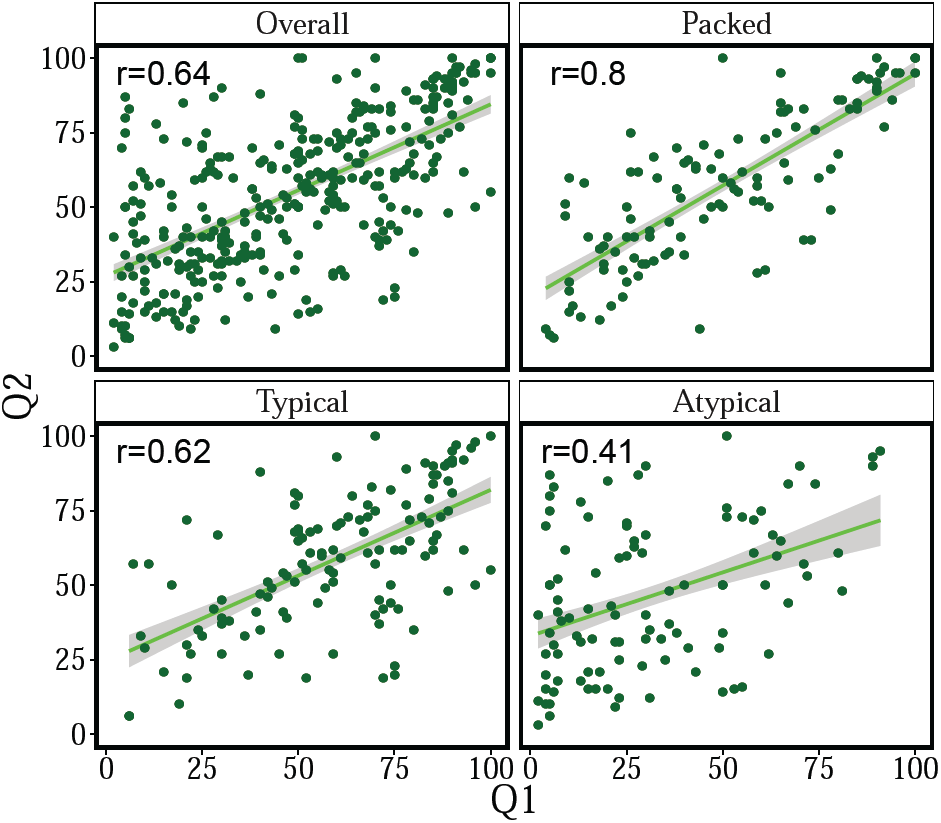
Experiment 1 trial-by-trial analyses: Relationship between aggregated *Q*1 and *Q*2 responses. Lines show the least-squares fit with standard error bands.

## Experiment 2

Experiment 1 showed strong evidence for reuse of inferential computations across queries when the evidence is fixed. Two questions naturally arise from this finding. First, how adaptive is amortization? Are samples reused promiscuously across queries (potentially leading to rampant memory-based biases), or is reuse sensitive to conditions where it is likely to be accurate? This is a delicate point, since it is impossible to know with certainty whether amortization is useful without knowing some properties of the problem (e.g., decomposability of the conditional distribution). Nonetheless, humans may be able to utilize heuristics for constructing amortization strategies whose errors can be corrected by additional experience or computation (Stuhlmüller et al., 2013). We address this question by manipulating the “amortizability” of *Q*1, in order to test the hypothesis that carry-over effects across queries will only occur under high amortizability conditions. We operationalize amortizability in terms of whether or not the hypothesis space queried by *Q*1 is a subset of the hypothesis space queried by *Q*2.

The second question concerns resource allocation. Theories of computational rationality argue that computations are selected to balance accuracy and cost (Lieder et al., 2012; Gershman et al., 2015). In the context of sampling, this means that fewer samples will be generated when cognitive resources are scarce. This hypothesis is consistent with the observation that subadditivity (Sprenger et al., 2011) and order effects (Thaker et al., 2017) are amplified under cognitive load. We pursue this question by manipulating cognitive load at both *Q*1 and *Q*2. As discussed in the Introduction, the different amortization schemes make different predictions for these manipulations (see Figure 1).

### Participants

80 participants (53 males, mean age=32.96, SD=11.56) were recruited from Amazon Mechanical Turk and received $0.5 as a basic participation fee and an additional bonus of $0.1 for every on time response as well as $0.1 for every correctly classified digit during cognitive load trials.

### Procedure

The procedure in Experiment 2 was largely the same as in Experiment 1, with the main difference being that participants had to remember a sequence of digits. On half of the trials the cognitive load manipulation occurred at *Q*1 and on half at *Q*2. The digit sequence was presented prior to the query, and then following their response to the query they were asked to make a same/different judgment about a probe sequence. Half of the probes were old and half were new.

To probe adaptive amortization, we added several *Q*2 queries to the list shown in Table 1. These queries were deemed less amortizable because they lack any of the letters queried in *Q*1 (for example, ’T or R’ instead of ‘C or R’ in the trial shown in the first row in Table1). In other words, these queries could not be decomposed and hence could not benefit from reuse. Half of the *Q*2 trials were randomly chosen to provide hypotheses with low amortizability.

### Results

As shown in Figure 5, we replicated and extended the results from Experiment 1, showing both subadditivity and superadditivity effects for *Q*1 that carried over to *Q*2. Analyzing only amortizable queries (averaging across load conditions), we found that probability judgments for *Q*1 were higher for unpacked-typical compared to packed (subadditivity; *t*(79) = 4.38, *p <* 0.001) and lower for unpacked-atypical compared to packed (superadditivity *t*(79) = −4.94, *p <* 0.001). These same effects occurred for *Q*2 (unpacked-typical vs. packed: *t*(79) = 2.44, *p <* 0.01; unpacked-atypical vs. packed: *t*(79) = −1.93, *p <* 0.05).

**Figure 5:**
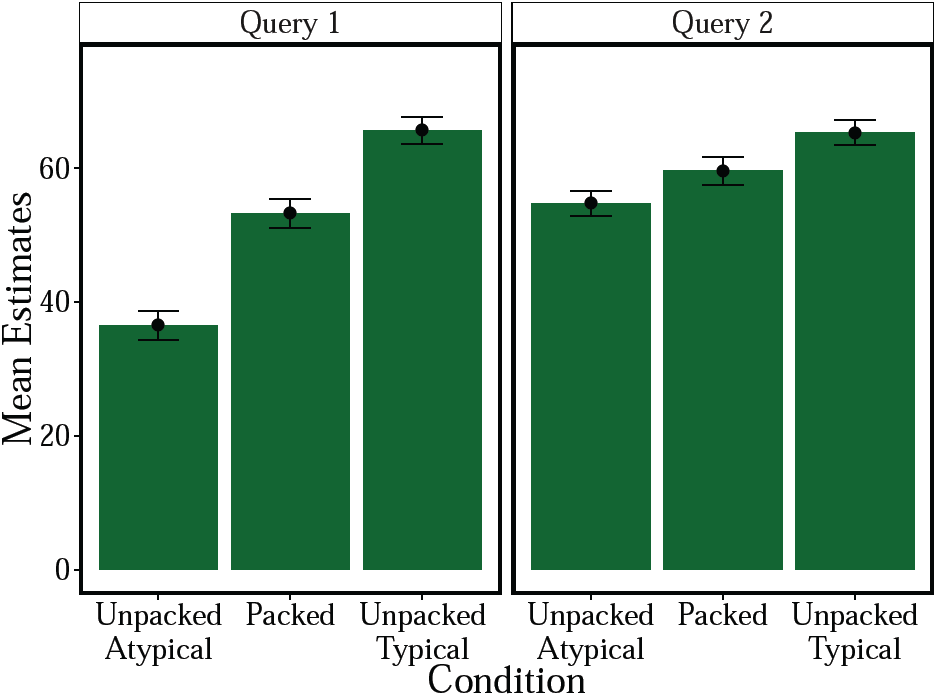
Experiment 2 results.

Mean probability estimates for *Q*2 by condition. Error bars represent the standard error of the mean.

As in Experiment 1, there was strong correlation between responses to *Q*1 and *Q*2 overall conditions (*r* = 0.44, *p <* 0.001), for the packed (*r* = 0.44, *p <* 0.001), the typically unpacked (*r* = 0.45, *p <* 0.001), as well as the atypically unpacked condition (*r* = 0.35, *p <* 0.01); see Figure 6. Moreover, *Q*1 and *Q*2 were also highly correlated within participants (mean *r* = 0.31, *p <* 0.01) and participants who showed higher subadditivity or superadditivty effects for *Q*1 also showed higher effects for *Q*2 overall (*r* = 0.31, *p <* 0.001), for the superadditive (*r* = 0.3, *p <* 0.001), and for the subadditive condition (*r* = 0.29, *p <* 0.001). This replicates the effects of amortization found in Experiment 1.

**Figure 6:**
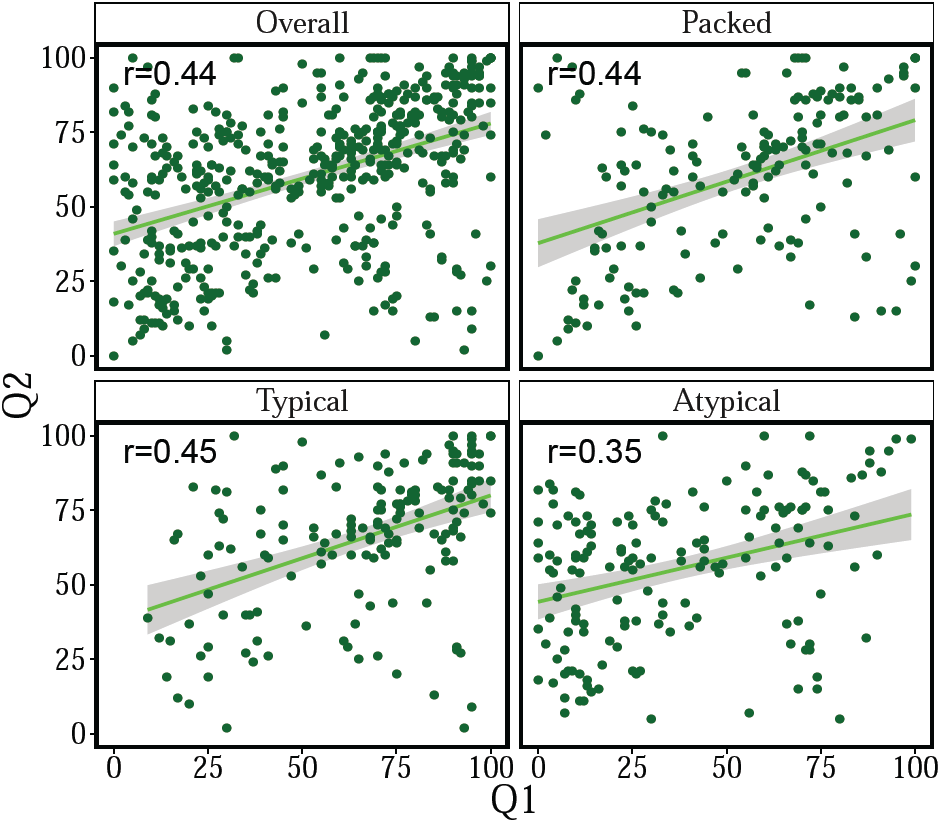
Experiment 2 trial-by-trial analyses: Relationship between aggregated *Q*1 and *Q*2 responses. Lines show the leastsquares fit with standard error bands.

Finally, we assessed how the carryover effects were modulated by cognitive load and amortizability. To highlight the effects more clearly, we calculated each participant’s mean response to all packed hypotheses for *Q*2 over all trials as a baseline measure. We then calculated the difference between each condition’s mean response and this mean packed response. This provides us with a measure of an average effect size within *Q*2-responses (how much each participant under- or overestimates different hypotheses as compared to an average packed hypothesis). Results are shown in Figure 7.

**Figure 7:**
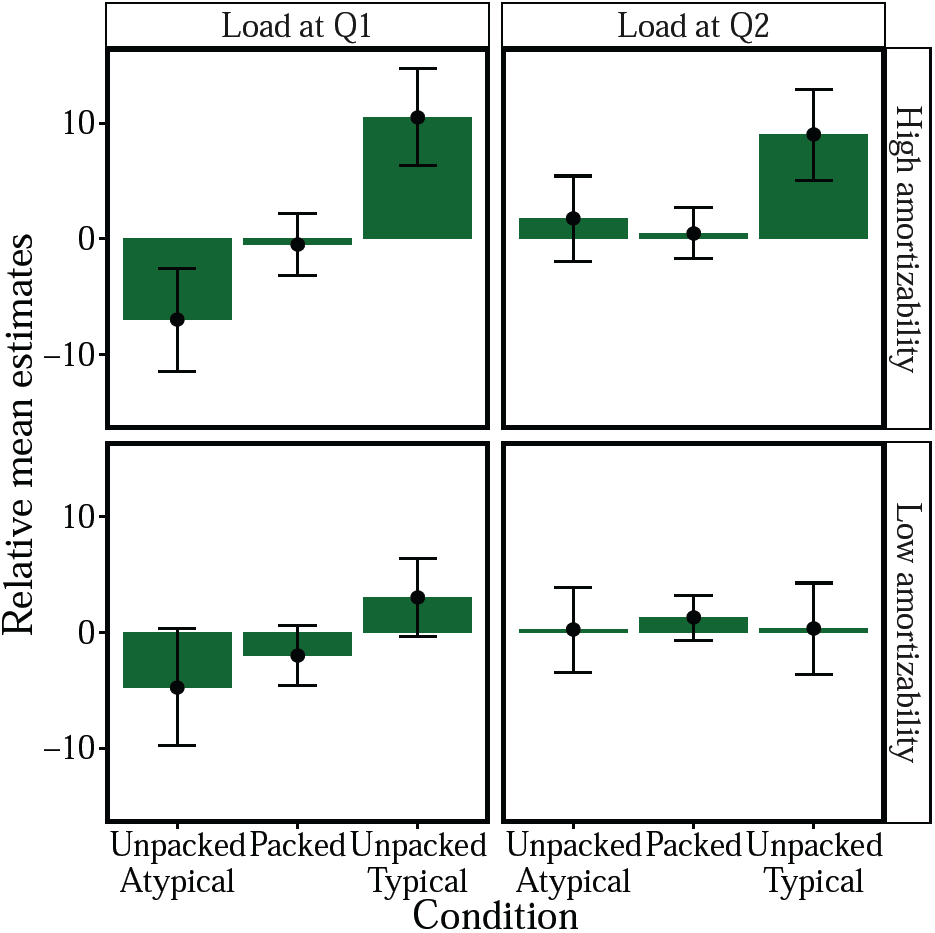
Experiment 2: differences between responses for each condition and an average packed baseline. Bars represent a mean within-participant absolute effect. Error bars represent the standard error of the mean.

If cognitive load occurred during *Q*2 and amortizability was low, none of the conditions produced an effect significantly different from 0 (all *p >* 0.5). If cognitive load occurred during *Q*2 and amortizability was high, only typically unpacked hypotheses produced an effect significantly higher than 0 (*t*(38) = 2.14, *p <* 0.05). If cognitive load occurred during *Q*1 and amortizability was low, again none of the conditions significantly differed from 0 (all *p >* 0.05). Crucially, if cognitive load occurred during *Q*1 and amortizability was high, both conditions showed the expected subadditive (*t*(38) = 4.18, *p <* 0.05) and superadditive (*t*(46) = −1.89, *p <* 0.05) effects. Moreover, calculating the average effect size for the different quadrants of Figure 7, the high amortizability-cognitive load at *Q*1-condition produced the highest overall effect (*d* = 0.8), followed by the high amortizability-cognitive load at *Q*2-condition (*d* = 0.56) and the low amortizability-cognitive load at *Q*1-condition (*d* = 0.42). The low amortizability-cognitive load at *Q*2-condition did not produce an effect higher than 0. Moreover, highly amortizable trials were more strongly correlated with responses during *Q*1 than trials with low amortizability (0.15 vs 0.41, *t*(157) = −2.28, *p <* 0.05).

Intriguingly, on trials with cognitive load at *Q*2, participants were on average more likely to answer the probe correctly for high amortizability trials compared to low amortizability trials (*t*(36) = 3.16, *p <* 0.05). This is another signature of amortization: participants are expected to have more resources to spare for the memory task at *Q*2 if the computations they did for *Q*1 are re-usable in answering *Q*2. This also indicates that these results cannot be explained by simply initializing the chain for *Q*2 where the chain for *Q*1 ended, which would not have affected computation time. Our results suggest that the transfer actually makes the second computation easier by re-using previous computations.

In summary, Experiment 2 replicates the effects found in Experiment 1 and the increased effect for the high amortizability condition provides further evidence that this effect is actually driven by amortization. These experiments also give us some insight into how amortization is implemented. Based on our simulations (Figure 1), we argue that the effect of cognitive load at *Q*1 on *Q*2 responses is more consistent with *summary amortization* than with *sample amortization*. These results suggest an active process of *Q*2 being expressed in terms of the results to *Q*1, when possible. This approach is more structured and less flexible than *sample amortization* but trades in this inference limitation for an increase in memory-efficiency and is thus consistent with beliefs about cost-efficient resource-rational inference strategies in humans.

## Discussion

In two experiments, we provided empirical support for amortized hypothesis generation. Participants not only exhibited subadditive and superadditive probability judgments in the same paradigm (replicating Dasgupta et al., 2016), but also carried over these effects to subsequent queries. Importantly, Experiment 2 demonstrated that such carry-over effects only occur when amortization can exploit shared structure across queries. Experiment 2 also demonstrated that cognitive load exerts its strongest effect when applied to the first query, suggesting (based on our simulations) that the carry-over effects are driven by some kind of summary-based amortization, whereby a summary statistic is computed from the samples and then reused to answer subsequent queries that can be expressed in terms of already completed calculations. This implies a structured amortization strategy, over one that reuses all old samples, and thus gives up some flexibility for memory-efficiency. Building on earlier results (Gershman & Goodman, 2014), our results support the existence of a sophisticated inference engine that adaptively exploits past computations. While reuse can introduce error, this error may be a natural consequence of a resource-bounded system that optimally balances accuracy and efficiency (Lieder et al., 2012; Vul et al., 2014; Griffiths et al., 2015; Gershman et al., 2015). The incorporation of reuse into a Monte Carlo sampling framework allows the inference engine to preserve asymptotic exactness while improving efficiency in the finite-sample regime.

Future studies could use similar methods to study amortization in other domains, such as in concept learning (Goodman et al., 2008) or reinforcement learning tasks (Daw et al., 2011). There is also a much larger space of more sophisticated amortization schemes (e.g., Stuhlmüller et al., 2013; Rezende et al., 2014; Paige & Wood, 2016) that we have not yet tried to test. Pinning down the computational details of amortization will be an important task for future work.

## Footnotes

1 Although more sophisticated amortization schemes have been developed in the machine learning literature (e.g., Stuhlmüller et al., 2013; Rezende et al., 2014; Paige & Wood, 2016), they are difficult to test experimentally in humans.

## References

Dasgupta, I., Schulz, E., & Gershman, S. J. (2016). Where do hypotheses come from? CBMM Memo 56.

Daw, N. D., Gershman, S. J., Seymour, B., Dayan, P., & Dolan, R. J. (2011). Model-based influences on humans’ choices and striatal prediction errors. Neuron, 69(6), 1204–1215.

Gershman, S. J., & Goodman, N. D. (2014). Amortized inference in probabilistic reasoning. In Proceedings of the 36th Annual Conference of the Cognitive Science Society, (pp. 517–522).

Gershman, S. J., Horvitz, E. J., & Tenenbaum, J. B. (2015). Computational rationality: A converging paradigm for intelligence in brains, minds, and machines. Science, 349(6245), 273–278.

Goodman, N. D., Tenenbaum, J. B., Feldman, J., & Griffiths, T. L. (2008). A rational analysis of rule-based concept learning. Cognitive Science, 32(1), 108–154.

Greene, M. R. (2013). Statistics of high-level scene context. Frontiers in Psychology, 4(1), 777.

Griffiths, T. L., Lieder, F., & Goodman, N. D. (2015). Rational use of cognitive resources: Levels of analysis between the computational and the algorithmic. Topics in Cognitive Science, 7, 217–229.

Griffiths, T. L., Vul, E., & Sanborn, A. N. (2012). Bridging levels of analysis for probabilistic models of cognition. Current Directions in Psychological Science, 21(4), 263–268.

Lieder, F., Griffiths, T. L., & Goodman, N. D. (2012). Burn-in, bias, and the rationality of anchoring. In Advances in Neural Information Processing Systems, (pp. 2690–2798).

MacKay, D. J. (2003). Information theory, inference and learning algorithms.

Paige, B., & Wood, F. (2016). Inference networks for sequential monte carlo in graphical models. In Proceedings of the 33rd International Conference on Machine Learning, vol. 48.

Rezende, D. J., Mohamed, S., & Wierstra, D. (2014). Stochastic backpropagation and approximate inference in deep generative models. In Proceedings of The 31st International Conference on Machine Learning, (pp. 1278–1286).

Sanborn, A. N., & Chater, N. (2016). Bayesian brains without probabilities. Trends in Cognitive Sciences, 20(12), 883–893.

Sloman, S., Rottenstreich, Y., Wisniewski, E., Hadjichristidis, C., & Fox, C. R. (2004). Typical versus atypical unpacking and superadditive probability judgment. Journal of Experimental Psychology: Learning, Memory, and Cognition, 30(3), 573–582.

Sprenger, A. M., Dougherty, M., Atkins, S. M., Franco-Watkins, A. M., Thomas, R., Lange, N., & Abbs, B. (2011). Implications of cognitive load for hypothesis generation and probability judgment. Frontiers in Psychology, 2, 129.

Stuhlmüller, A., Taylor, J., & Goodman, N. D. (2013). Learning stochastic inverses. In Advances in Neural Information Processing Systems, (pp. 3048–3056).

Thaker, P., Tenenbaum, J. B., & Gershman, S. J. (2017). Online learning of symbolic concepts. Journal of Mathematical Psychology.

Tversky, A., & Koehler, D. J. (1994). Support theory: a nonextensional representation of subjective probability. Psychological Review, 101, 547–567.

Vul, E., Goodman, N., Griffiths, T. L., & Tenenbaum, J. B. (2014). One and done? optimal decisions from very few samples. Cognitive Science, 38(4), 599–637.

